# Prefrontally Modulated Vagal Tone Inhibits Inflammatory Responses to Prevent Telomere Damage in Healthy Participants

**DOI:** 10.1101/2022.02.17.480574

**Authors:** Torvald F. Ask, Stefan Sütterlin

## Abstract

**Background:** Accumulated senescent cells are proposed to be one of the main drivers of age-related pathology through disruption of tissue structure and function. We recently proposed the Neuro-Immuno-Senescence Integrative Model (NISIM; Ask et al., 2018) which relates prefrontally modulated vagal tone and subsequent balance between vagal and sympathetic input to the spleen to inflammatory responses leading to generation of reactive oxygen species and oxidative telomere damage. The NISIM is based on converging evidence and argues for the existence of a prefrontal cortex-autonomic nervous system-spleen (PAS) axis, suggesting that the inflammation that induces reactive oxygen species-generation is downstream of this axis.

**Aim:** In this study, we aim to assess inflammation as a mediator in the relationship between prefrontally modulated vagal tone and leukocyte telomere length to test the hypothesis that PAS axis dysregulation accelerates cellular aging. We also assess the relationship between a recently proposed index of vagal immunomodulation (vagal tone/inflammation ratio; NIM index; Gidron et al., 2018) and telomere length, and compare results between the NIM index and vagal tone as predictors of telomere length.

**Methods:** This study uses participant data from a large nationally representative longitudinal study since 1974 with a total of 45,000 Norwegian residents so far. A sub-sample of 1372 participants from which vagal tone, C Reactive Protein, and leukocyte telomere length could be obtained were included in the study. Relationships were analyzed with hierarchical multiple linear regression using either vagal tone and C Reactive Protein or the NIM index to predict telomere length. Sleeping problems, tobacco use status, alcohol use status, time since last meal, and symptoms of depression were included as control variables.

**Results:** In the mediation analysis, vagal tone was a significant positive predictor of telomere length, while C Reactive Protein was a significant negative predictor of telomere length. This relationship remained significant when individually controlling for some but not all confounding variables. The NIM index was a significant positive predictor of telomere length. This relationship remained significant when controlling for all confounding variables except one. In a reduced dataset excluding all participants where confounders were present, the NIM index remained a significant predictor of telomere length.

**Conclusion:** This is the first study suggesting that PAS axis activity is associated with telomere length thus supporting the NISIM. Results indicate that the NIM index is a more sensitive indicator of PAS axis activity than vagal tone and C Reactive Protein in isolation. Clinical relevance and suggestions for future research are discussed.

## 1 Introduction

In this paper, we test the previously postulated hypothesis that inflammation mediates the relationship between vagally mediated heart rate variability (vmHRV) and telomere length (Ask, Lugo, & Sütterlin, 2018) to elucidate the mechanisms linking individual differences in psychophysiological stress regulation capacity to aging and age-related pathophysiology. We also test whether a recently proposed index for vagal immunomodulation (Gidron et al., 2018) is associated with telomere length and we discuss it with respect to the mediation analysis. The shift towards considering aging as a disease has prompted increasing efforts to understand its many underlying mechanisms and their subsequent role in degenerating pathologies (e.g. Campisi, 2000, 2013; Kane & Sinclair, 2019). These efforts converge on the ambition that if humans can fully understand the physiological drivers of aging then perhaps medical- and behavioral interventions can be developed such that age-related pathologies can be eradicated altogether.

Aging entails a progressive loss of physiological integrity resulting in impaired function and increased vulnerability to death, with hallmarks being genomic instability, telomere attrition, epigenetic alterations, loss of proteostasis, deregulated nutrient-sensing, mitochondrial dysfunction, stem cell exhaustion, cellular senescence, and altered intercellular communication (Figure 1, a; López-Otín et al., 2013). These hallmarks can be divided into three categories: primary hallmarks (causes of damage), antagonistic hallmarks (responses to damage), and integrative hallmarks (culprits of the [aging] phenotype).

**Figure 1.**
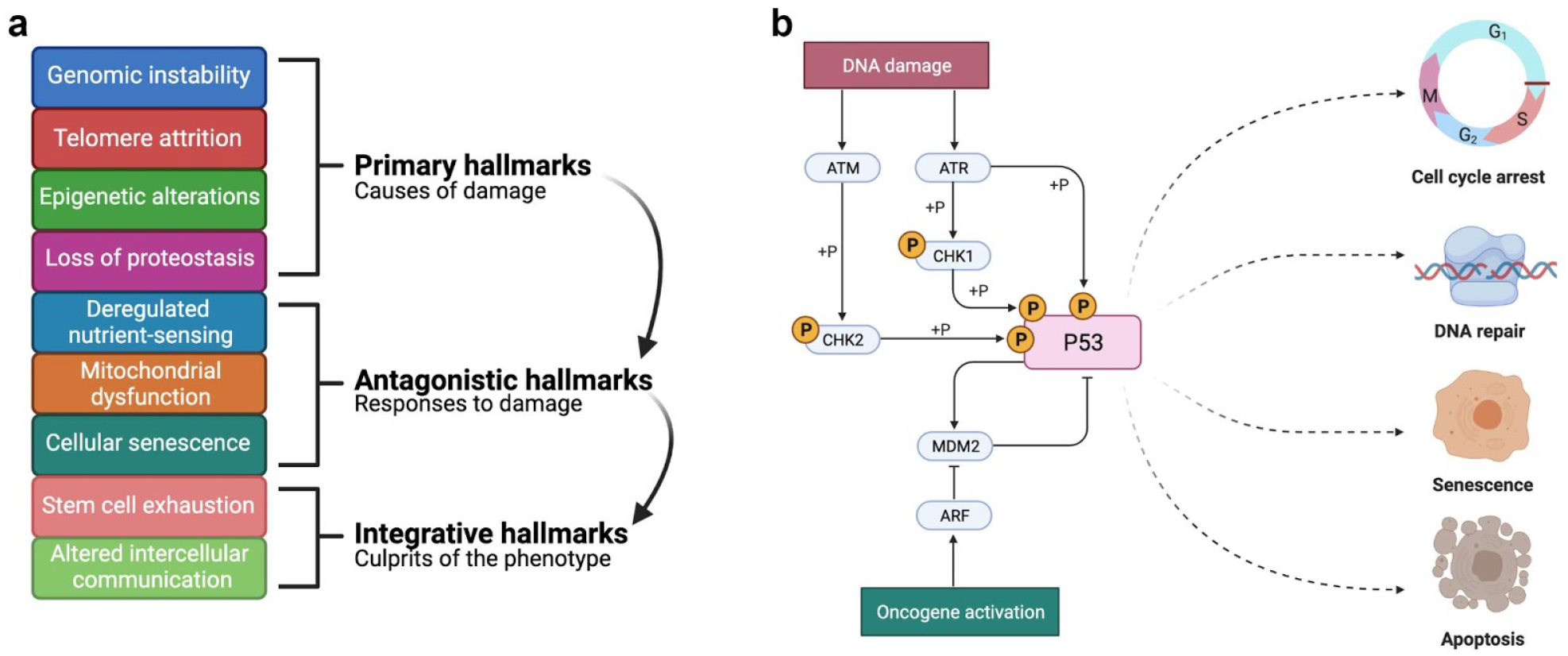
The hallmarks of aging and the DNA damage response. **a** The three categories of the hallmarks of aging. Modified with permission from López-Otín et al. (2013). **b** The P53-mediated DNA damage response pathway. ATM = ATM Serine/threonine-protein kinase. ATR = ATR Serine/threonine-protein kinase. CHK1 = Checkpoint kinase 1. CHK2 = Checkpoint kinase 2. P53 = protein 53. MDM2 = Mouse double minute 2. ARF = Alternative reading frame. Created with BioRender.com.

**Figure 2.**
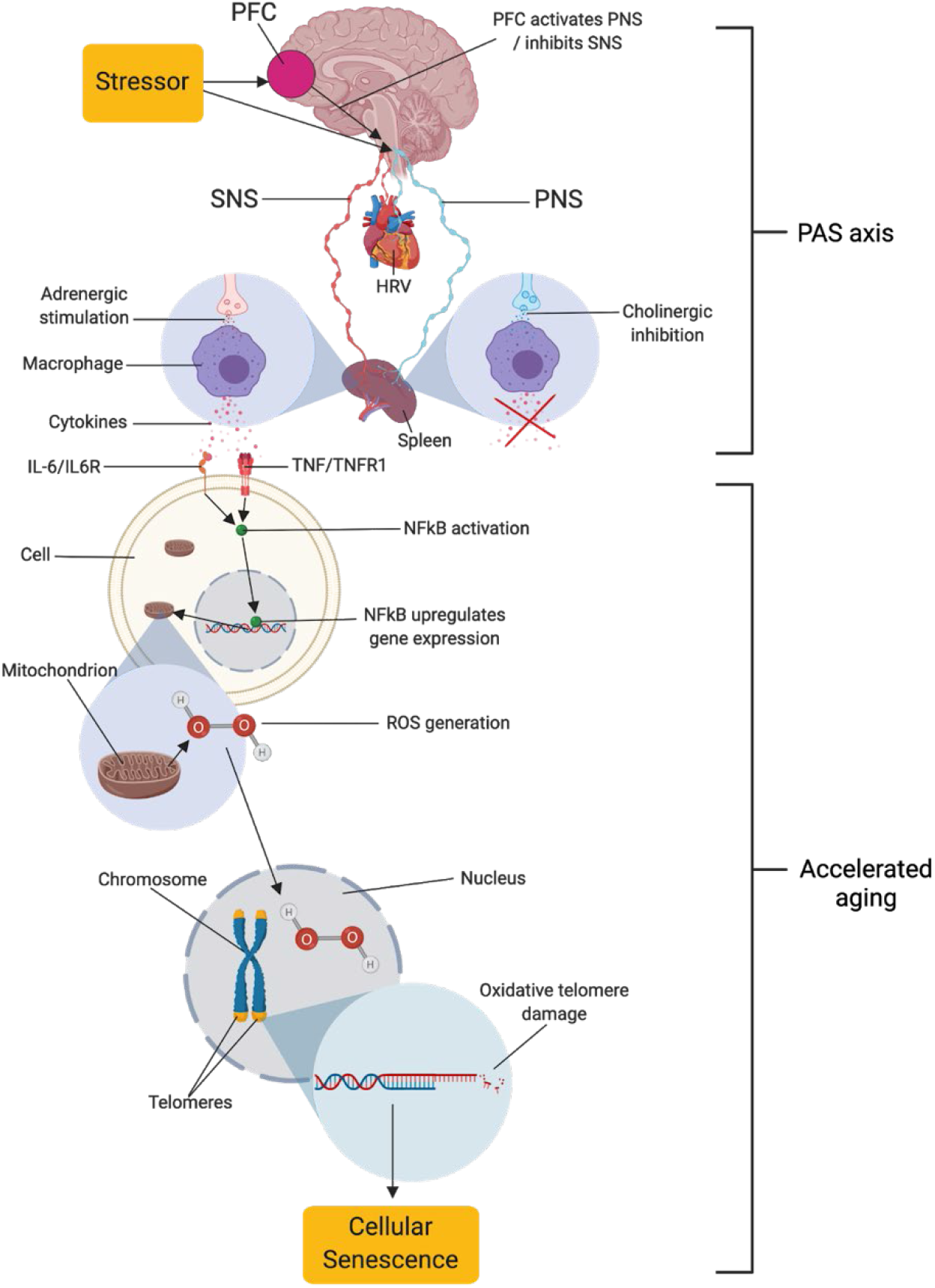
The neuro-immuno-senescence integrative model (NISIM). The NISIM propose that the link between psychological stress and telomere length starts with (1) prefrontal capacity for increasing activity in the vagus nerve, (2) vagal input to the spleen modulating the inflammatory response (3), subsequent inflammatory load increasing Nuclear Factor Kappa B (NFkB) activity and reactive oxygen species generation, and (4) the resulting oxidative damage to telomeres (Ask et al., 2018). PFC = Prefrontal cortex. SNS = Sympathetic nervous system. PNS = Parasympathetic nervous system. HRV = Heart rate variability. PAS axis = Prefrontal cortex-autonomic nervous system-spleen axis. IL-6 = Interleukin 6. IL6R = IL6 receptor. TNF = Tumor Necrosis Factor. TNFR1 = TNF receptor 1. ROS = Reactive oxygen species. HOOH = Hydrogen peroxide. Figure created with BioRender.com.

Cellular senescence, also known as cellular aging, is secondary to some of these hallmarks, most notably genomic instability, telomere attrition, and mitochondrial dysfunction (Campisi, 2000, 2013). Following DNA damage, a network cascade of proteins is activated as part of the DNA damage response (DDR), leading to the phosphorylation of P53. Whether DNA-damage induced P53 activation will result in cellular senescence or another response such as DNA repair or apoptosis depends on the persistence of the DDR signal and the cellular stressor (Campisi, 2000, 2013; Figure 1, b). Due to being resistant to apoptosis, senescent cells accumulate with age. Some senescent cells start secreting a wide range of ligands including inflammatory molecules and growth factors that over time promote disruption in tissue structure and function, thus altering intercellular communication (Campisi, 2000, 2013; Tchkonia et al., 2013). This is known as the senescence-associated secretory phenotype and is a suggested driver of age-related pathologies such as neurodegenerative diseases (Baker et al., 2011; Chinta et al., 2015) and cancer (Bissel et al., 2005; Rao and Jackson, 2016). Individual anato-physiological factors driving the antecedents to cellular senescence are still poorly understood.

### 1.1 The Neuro-Immuno-Senescence Integrative Model (NISIM) on the relationship between prefrontally modulated vagal tone and cellular senescence

In a seminal paper, Epel and colleagues (2004) made the pioneering discovery that psychological stress is related to shorter telomere length, an antecedent to cellular senescence (Campisi, 2000, 2013). While being a critical contribution to our understanding of how psychological stress relates to ill-health, the specific mechanisms relating these phenomena have remained rather elusive for the remaining decade-plus despite considerable scientific inquiry (e.g. Epel et al., 2009; Morey et al., 2015). In an effort to bridge this gap, we recently proposed a model (Figure 1) outlining specific anato-physiological mechanisms relating psychological stress regulation to cellular senescence, and subsequently, age-related pathology (Ask, Lugo, & Sütterlin, 2018). The model is largely based on the prefrontal component of the central-autonomic network, the immuno reflex (or cholinergic anti-inflammatory pathway; Tracey, 2002), and the molecular cell biology related to generation of reactive oxygen species (ROS; e.g. Schreck & Bauerle, 1994). In this model, termed the Neuro-Immuno-Senescence Integrative Model (NISIM; Ask et al., 2018), the prefrontal capacity for modulation of vagal activation, quantified as vmHRV (Thayer & Lane, 2000; Thayer et al., 2012) during exposure to psychological stress, consequently affect the balance of vagal and sympathetic input to the spleen (Tracey, 2002). Low capacity for prefrontal vagal modulation translates to low vmHRV (Thayer & Lane, 2000), indicating reduced vagal input to the spleen and increased inflammation (Williams et al., 2019). Increased inflammatory load, such as increased levels of Interleukin-6 (IL-6), C-Reactive Protein (CRP), and Tumor Necrosis Factor α (TNF-α) is negatively associated with telomere length (Fitzpatrick et al., 2006; Masi et al., 2012; O’Donovan et al., 2011; Wong et al., 2014), while indicators of vagal tone such as respiratory sinus arrhythmia (Kroenke, 2011) and vmHRV is positively associated with telomere length (Perseguini et al., 2015; Streltsova et al., 2017; Woody et al., 2017; Zalli et al., 2013). Together, the outlined above mechanisms are how the NISIM relates psychological stress to cellular senescence in a dynamic framework that accounts for inter-individual differences in the expression of age-related phenotypes.

It is important to note that the mechanisms outlined by the NISIM are not suggested to be the main drivers of cellular senescence. The NISIM should rather be viewed as detailing the neuroimmunological mechanisms underlying individual differences in small but cumulative antecedents to cellular senescence that accelerate the transition from health to age-related pathology (Ask et al., 2018). Higher vmHRV has been linked to increased life-span (Piccirillo et al., 1998; Shimizu et al., 2002), while both lower vmHRV and proxies of low vmHRV predict all-cause mortality (Tsuji et al., 1994), cancer onset (Kennedy et al., 2017), Parkinson disease onset (Alonso et al., 2015), Alzheimer’s disease onset (Wang et al., 2012), is associated with cognitive decline (Kim et al., 2007; Al Hazzouri et al., 2017), and degree of cognitive impairment in Alzheimer’s (Santos et al., 2017; Zulli et al., 2006). The NISIM proposes that these relationships can in part be attributed to low or reduced prefrontal modulation of vagal activity resulting in inflammatory responses that accelerate cellular aging (Ask et al., 2018). Although numerous evidence converges on this interpretation, studies simultaneously assessing indicators of prefrontally mediated vagal tone, markers of inflammation, and their relationships with antecedents to cellular senescence has yet to be conducted. Analyzing these relationships are the current aim of the present study.

### 1.2 The prefrontal cortex-autonomic nervous system-spleen (PAS) axis and its role in neuroimmunomodulation

Senescent cells are adaptive in a young organism but can be detrimental to the health of older organisms in which they tend to accumulate (Campisi, 2000). The idea that the same biological function (e.g. the genes regulating cellular senescence) can produce two seemingly opposing phenotypes (health and disease) is called antagonistic pleiotropy (Williams, 1957). The evolutionary explanation for why cellular senescence protects young organisms but promotes age-related pathology in older organisms has been described elsewhere (e.g. Ask et al., 2018; Campisi, 2000). In short, the evolutionary account attributes the opposing phenotypes of healthy-young and sick-old to the fact that cellular senescence evolved as an adaptive response to cellular stress in organisms that had a much shorter life-span than humans have today. The same evolutionary principle can be attributed to inflammation. Psychological stress and arousal is associated with increased expression of inflammatory markers (Maydych, 2019). In the wild, organismal stress usually occurs when life is in danger and increased inflammatory expression prepares the organism to battle infections that might occur in wounds from fighting other organisms (Semple et al., 2002). As humans live longer due to improved standards of living, stress-induced inflammatory responses that protect the young organism from infection become a driver of age-related pathology in older organisms through induction of chronic low-grade inflammation (Liu et al., 2017).

Due to the capacity of the human brain to represent and solve problems cognitively while being physically and temporally removed from the source of the problem, humans can mentally represent and maintain psychological stressors through perseverative cognitions (Brosschot et al., 2005). Perseverative cognitions are associated with increased levels of inflammation (e.g. Moriarity et al., 2019). This inflammation-associated problem-solving ability might be helpful if it is in anticipation of an actual and imminent threat (e.g. a medical doctor knowing they are about to enter an environment with high levels of infection) by subtly boosting inflammation prior to exposure. If an individual is not capable of disengaging from perseverative thought in the absence of a stressful context, it can be a health risk by sustaining responses that contribute to the chronic low-grade inflammation associated with aging (Moriarity et al., 2019). Thus, psychological stress can contribute to inflammation in humans either as a result of direct exposure to the stressor or through the evolved ability for perseverative mental representation of stressors.

Stressors, and the resulting levels of arousal can induce inflammatory responses in several ways, one of which includes pathways via the autonomic nervous system (ANS) fibers descending into lymphoid tissues such as the spleen (Felten & Felten, 1994; Rosas-Ballina et al., 2008). Sympathetic nerve fibers can induce inflammation by transmitting adrenergic ligands onto α-subtype adrenoceptors on leukocytes in the spleen (Madden et al., 1995; Vizi, 1998) while vagal nerve fibers can suppress inflammation (Tracey 2002, 2007) through the transmission of cholinergic ligands onto the α7 nicotinic acetylcholine receptors on leukocytes (Huston et al., 2006; Wang H et al., 2003; Wang, 2004). The cholinergic modulation of inflammation is mediated by the splenic nerve (Rosas-Ballina et al., 2008). This makes the spleen a neuroimmune interface that allows the nervous system to adapt inflammatory responses to situational stressors (Noble et al., 2018; Rosas-Ballina et al., 2008).

Through its expansion, the prefrontal cortex (PFC) is arguably the most recently evolved structure in the human brain and is involved in executive functioning and in the adaptive regulation of the physiological, emotional, cognitive, and behavioral responses to acute and chronic stress (Appelhans & Luecken, 2006; Gross et al., 1998). The PFC achieves this in part by modulating activity in the ANS through its connections with the amygdala and pre-autonomic cell groups in the hypothalamus, periaqueductal grey, and brainstem (Golkar et al., 2012; Seeley et al., 2007). To reduce arousal, the PFC can either increase activity in the vagus nerve or inhibit activity in the sympathetic nervous system. Higher prefrontal activity is reflected in higher vmHRV (Jennings et al., 2017; Nikolin et al., 2017; Thayer et al., 2012), an indicator of prefrontally modulated vagal tone, and higher vmHRV predicts better regulation of physiological arousal during prolonged stressors (Hildebrandt et al., 2016). Due to its ability to modulate ANS activity, the PFC is arguably a central component in adapting the ANS-modulated inflammatory response to situational needs. Consequently, lower prefrontal capacity for ANS modulation will result in inappropriate levels of inflammation in response to stress. Recent research showed that there was an inverse relationship between perseverative cognitions, PFC-associated executive functioning (e.g. Cropley & Collins, 2020) and vmHRV (Cropley et al., 2017), linking PFC activity to inflammation. Further evidence for the relationship between PFC function and inflammation was confirmed in a meta-analysis linking the activity in several prefrontal structures with the peripheral expression of inflammatory markers (Kraynak et al. 2018). Together, the interplay between these anatomical components constitute a prefrontal cortex-autonomic nervous system-spleen (PAS) axis, suggesting that an individual’s capacity for regulating stress responses through prefrontal modulation of vagus nerve activity has downstream effects on inflammation. To the best of our knowledge, this is the first paper conceptualizing these relationships as an axis.

The relationship between vagal tone and inflammatory markers has been reviewed in studies using healthy human participants (Ask et al., 2018) and participants with cardiovascular diseases (Haensel et al., 2008; Papaioannou et al., 2013). Although indicative of a relationship, evidence remained rather inconclusive. In a recent meta-analysis (Williams et al., 2019), the relationship between HRV and inflammatory marker-expression was revisited. The meta-analysis included data on several HRV indices and markers of inflammation, including IL-1, IL-2, IL-4, IL-6, and IL-10, CRP, leukocyte (white blood cell; WBC) count, Fibrinogen, Interferon-gamma (IFN-γ) and TNF-α. The high-frequency component of HRV (HFHRV) showed the strongest association to inflammatory markers of all vmHRV indices, with CRP and IL-6 showing the most reliable relationships with HFHRV (Williams et al., 2019). No significant associations were found between time domain measures of vmHRV (the Root Means Square of Successive Differences; RMSSD) and IL-6. An explanation for this discrepancy may be attributed to that RMSSD might be influenced by sympathetic activation (Berntson et al., 2005). Vagal (Huston et al., 2006; Wang H et al., 2003) and sympathetic (Madden et al., 1995; Vizi et al., 1998) inputs to the spleen have opposing effects on inflammatory marker expression. Moreover, IL-6 appears to have a stronger relationship with HFHRV than CRP (Williams et al., 2019), possibly due to CRP expression being downstream of IL-6 expression (Boras et al., 2014) as it is mainly synthesized through IL-6-dependent hepatic biosynthesis (Baumeister et al., 2016; Pradhan et al., 2001). No significant associations were found between TNF-α and vmHRV, although the authors note that the sample sizes for studies assessing these relationships were small (Williams et al., 2019). The authors concluded that there was evidence for a vagal modulation of inflammation in humans, providing statistical support for a PAS axis. The NISIM argues that the PAS axis is the common denominator in both how individual differences in stress regulation capacity translates to inflammation-related pathology as well as explaining why they are subtle and accumulate over time (Ask et al., 2018).

### 1.3 PAS axis dysregulation and implications for the NISIM

The NISIM suggests that a dysregulated PAS axis will result in increased inflammation-induced levels of reactive oxygen species (ROS), which in turn will accelerate ROS-induced telomere shortening resulting in cellular senescence (Ask et al., 2018). Telomeres are DNA-protein complexes consisting of base pairs of TTAGGG repeats that cap the ends of linear chromosomes to protect them from end-to-end fusion by DNA-repair processes and degradation (Blackburn, 1991; Verdun & Karlseder, 2006). When telomeres become dysfunctional, either through oxidative DNA damage or mitosis-associated telomere attrition, the DDR response is activated and if persisting will result in cellular senescence (Campisi, 2013; Coluzzi et al., 2019). Shorter telomeres have been reported to increase risk for Alzheimer’s (Thomas et al., 2008). Whether shorter telomeres are a risk for cancer development might depend on cancer type (Zhu et al., 2016), although telomere dysfunction can occur irrespective of telomere length (Victorelli & Passos, 2017). An increasing number of studies link inflammation to telomere length. A two-year longitudinal study found negative associations between CRP and telomere length (Wong et al., 2014) and genetically elevated CRP levels were found to be inversely associated with telomere length (Rode et al., 2014). Interestingly, an inverse relationship between CRP levels and telomere length was found among depressed young men, suggesting a relationship between emotion dysregulation and telomere attrition (Shin et al., 2019). Similarly, inverse relationships have also been found between telomere length and vmHRV (Perseguini et al., 2015; Streltsova et al., 2017; Woody et al., 2017; Zalli et al., 2013; but see also Epel et al., 2006).

While these data indicate that a dysregulated PAS axis contributes to the antecedents to cellular senescence, more direct measures of PAS axis activity with respect to telomere length are lacking. One method of approximating this problem would be to assess inflammation as a mediator in the relationship between vagal tone and telomere length.

### 1.4 The neuroimmunomodulation (NIM) index as an indicator of PAS axis activity

It is known that distressful psychological states associated with emotional dysregulation and lower PFC activity such as depression and loneliness are related to cancer progression and survival (Pinquart & Duberstein, 2010; Rico-Uribe et al., 2018). A series of studies associating vagal tone with cancer survival provided some of the first evidence suggesting that the ability of the PFC to modulate vagal activity and the resulting modulation of the immune system could be a manner of life and death in clinical conditions (Chiang et al., 2013; De Couck et al., 2013; Mouton et al., 2010, 2011, 2012). As vmHRV alone does not predict all outcomes (Gidron et al., 2005, 2018), and because it was found that CRP statistically mediates the relationship between vmHRV and survival in patients with pancreatic cancer (De Couck et al., 2015), there has been a need for a biomarker that can serve as a proxy relating vagal activity to modulation of the immune response. In absence of an index for this relationship, Gidron et al. (2018) proposed the neuroimmunomodulation (NIM) index; a simple numerical ratio (vmHRV/CRP) indexing vagal immunomodulation of inflammation. They found that the NIM index had a protective relative risk among pancreatic and non-small cell lung cancer patients and that non-small cell lung cancer patients with higher NIM index survived longer than those with lower NIM index scores. The authors suggested that this novel index could serve as an independent prognostic biomarker for cancer (Gidron et al., 2018). Inflammatory markers do not necessarily reflect vagal activity, nor does vagal activity necessarily reflect inflammation. Thus, the NIM index could potentially be a more sensitive measure of PAS axis activity than any of the measures in isolation. How NIM index scores, and by extent PAS axis activity, relate to antecedents of cellular senescence such as telomere length has yet to be explored.

### 1.5 Aim

The NISIM (Ask et al., 2018) proposes that for psychological effects on telomere length, prefrontal cortical influences on the vagus nerve modulate the inflammatory response with reduced telomere length being downstream of inflammation. Thus, the main aim of this paper is to test the hypothesis that inflammation negatively mediates the relationship between vagal tone and telomere length. The secondary aim is to assess whether the NIM index (Gidron et al., 2018) could serve as a useful index for the PAS axis in the NISIM by assessing whether the NIM index is a positive predictor of telomere length, and assess its sensitivity by comparing it to the mediation analysis. Due to the scarcity of existing data, the tertiary aim of this study is to replicate previous findings relating vagal tone to telomere length.

## 2 Methods

### 2.1 Ethics statement

This study uses participant data from the sixth survey of the Tromsø Study (Tromsø6; Eggen et al., 2013) which was collected from 2007 to 2008. Seven surveys have been carried out so far. Tromsø6 was approved by the Data Inspectorate of Norway and the Regional Committee of Medical and Health Research Ethics, North Norway. Tromsø6 complies with the Declaration of Helsinki, International Ethical Guidelines for Biomedical Research Involving Human Subjects and the International Guidelines for Ethical Review of Epidemiological Studies. Participation was voluntary and each subject gave written informed consent prior to participation.

### 2.2 Participants

The Tromsø Study cohort consists of people living in the municipality of Tromsø, Norway, situated at 69° North. Among the 70,000 people living in Tromsø when Tromsø6 was carried out, approximately 60,000 were living in the city-like town-center, while the remaining 10,000 were spread throughout the whole municipality (2558 km^2^). Tromsø is a center of education, research, administration and fishing related activities. It has a growing population that is predominantly Caucasians of Norwegian origin, but also includes a Sami minority. Tromsø may be considered as representative of a Northern European, white, urban population. A total of 12984 (male = 6054) participants aged 30-87 years (M = 57.5, SD = 12.7) enrolled in Tromsø6. Of these, 9946 participated in a cold-pressor test (described in Schistad et al., 2017) where blood pressure (BP) was measured for inter-beat-interval (IBI) extraction and vmHRV quantification. A total of 1372 participants from whom both telomere length and vmHRV measures could be obtained was included in the analysis. Characteristics of the study sample can be found in Table 1.

**Table 1.**
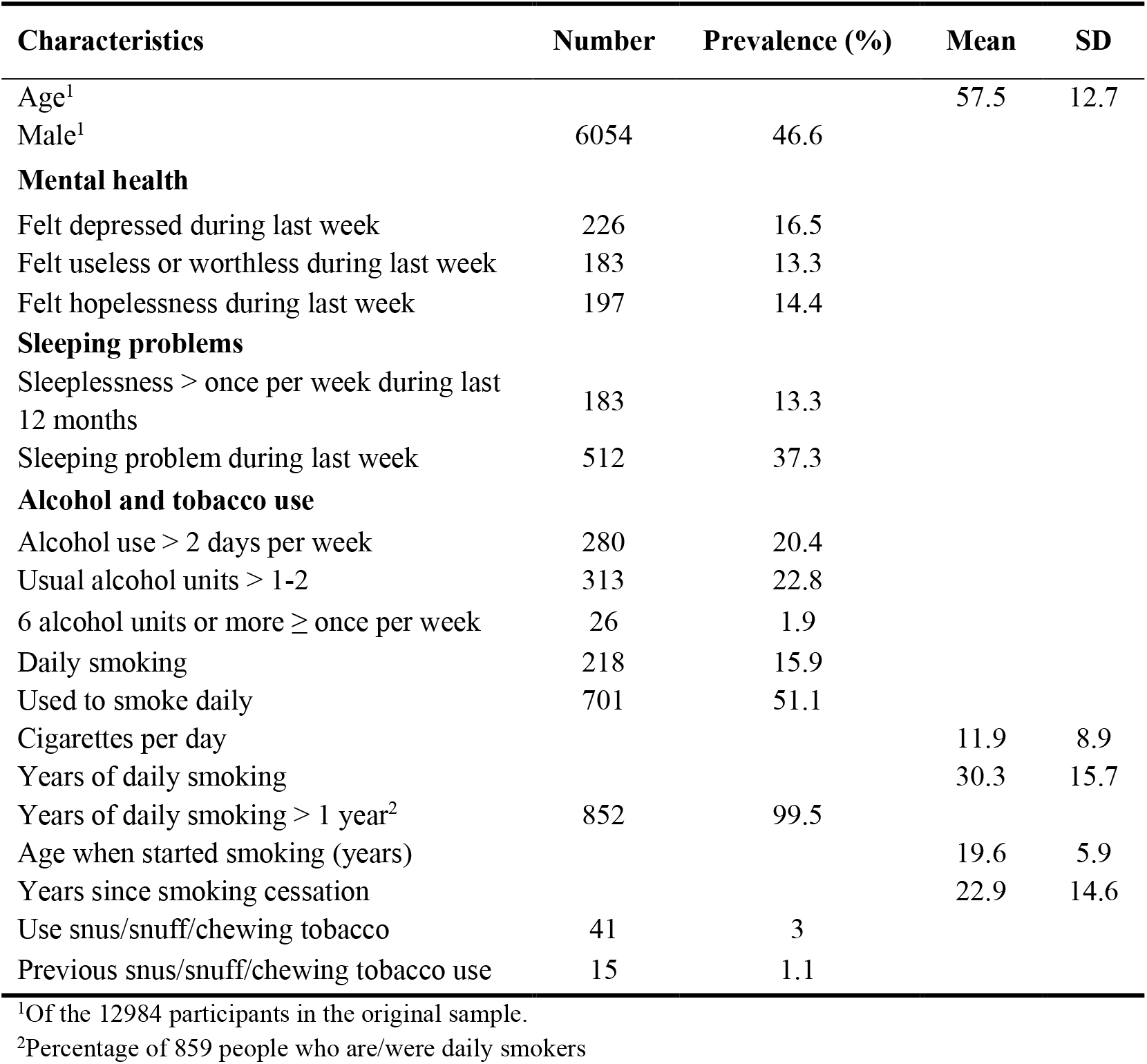
Study sample characteristics (N = 1372)

| Characteristics | Number | Prevalence (%) | Mean | SD |
| --- | --- | --- | --- | --- |
| Age<sup>1</sup> |  |  | 57.5 | 12.7 |
| Male<sup>1</sup> | 6054 | 46.6 |  |  |
| **Mental health** |  |  |  |  |
| Felt depressed during last week | 226 | 16.5 |  |  |
| Felt useless or worthless during last week | 183 | 13.3 |  |  |
| Felt hopelessness during last week | 197 | 14.4 |  |  |
| **Sleeping problems** |  |  |  |  |
| Sleeplessness > once per week during last 12 months | 183 | 13.3 |  |  |
| Sleeping problem during last week | 512 | 37.3 |  |  |
| **Alcohol and tobacco use** |  |  |  |  |
| Alcohol use > 2 days per week | 280 | 20.4 |  |  |
| Usual alcohol units > 1-2 | 313 | 22.8 |  |  |
| 6 alcohol units or more ≥ once per week | 26 | 1.9 |  |  |
| Daily smoking | 218 | 15.9 |  |  |
| Used to smoke daily | 701 | 51.1 |  |  |
| Cigarettes per day |  |  | 11.9 | 8.9 |
| Years of daily smoking |  |  | 30.3 | 15.7 |
| Years of daily smoking > 1 year<sup>2</sup> | 852 | 99.5 |  |  |
| Age when started smoking (years) |  |  | 19.6 | 5.9 |
| Years since smoking cessation |  |  | 22.9 | 14.6 |
| Use snus/snuff/chewing tobacco | 41 | 3 |  |  |
| Previous snus/snuff/chewing tobacco use | 15 | 1.1 |  |  |

<sup>1</sup>Of the 12984 participants in the original sample.
<sup>2</sup>Percentage of 859 people who are/were daily smokers

### 2.3 Vagally mediated heart rate variability (vmHRV)

#### 2.3.1 Recording, data reduction and analysis

BP was measured noninvasively for 30 seconds using a finometer device (Finometer, Finapres Medical System, Amsterdam, Netherlands) where the sensor was placed on the participant’s finger. The original dataset consisted of 9946 recordings of IBI over a mean duration of 310 samples from which heart periods were extracted, artifacts were corrected and time-domain and frequency-domain HRV parameters were derived. For pre-processing, artifact correction, and data formatting, Matlab R2015 (MATLAB, 2015) was used. For the calculation of all time-domain and frequency-domain HRV parameters, the RHRV package (Martínez et al., 2017) in the R language (R Core Team, 2020) was employed.

#### 2.3.3 Artifact correction

Artifacts was removed using a distribution-based algorithm that identifies artifacts as statistical outliers within a moving window and replaces them by linear interpolation (Deegan et al., 2007a, 2007b). The total distribution of IBI values (after the artifact correction) from all subjects was inspected in a histogram which revealed a small portion of values outside of the physiological range (i.e. IBI > 3s). A low and high cut-off limit for IBI observations of physiological origin was set to 0.3s and 2.5s, respectively. All recordings having at least one observation outside this limit were omitted (n = 378; 3.8% of the total of 9946).

#### 3.5.4 Calculation of HRV parameters in RHRV

Data files of the IBI for each recording were generated using a Matlab script then loaded into the RHRV script for HRV calculation. RHRV includes artifact removal by eliminating outliers or spurious points in the time-series with unacceptable physiological values. This method was compared to the Deegan et al. (2007a, 2007b) method by calculating all time-domain and frequency-domain parameters using both the raw and corrected (Deegan et al., 2007a, 2007b) IBI data. There was a significant disagreement between the two. Thus, the artifact-corrected data were used as input for the HRV calculations in RHRV.

#### 2.3.5 Time domain parameters

Due to the short data length of our recordings (30s), the triangular interpolation of NN histogram and HRV index parameters was calculated using the shape of the histogram of the RR intervals. To assess how bin width selection affected recordings, histograms of the default (8ms), lower (4ms) and higher (12ms) bin widths were compared for 25 random recordings. Plotted on top of each other, the 4ms bin gave a too sharp shape, while the 12 ms bin width gave a too blurry representation of the shape. Thus, the default setting of 7.8125 ms was used.

#### 2.3.6 Frequency domain parameters

For calculation of the frequency-domain parameters, Fourier-based signal power calculation in the LF, MF and HF bands were used. The whole recording duration is used as the window size in this calculation, thus there is no other setting for the Fourier method. The signal power was calculated in three bands (Task Force, 1996): LF (0.04-0.15 Hz), MF (0.07-0.15 Hz) and HF (0.15-0.4 Hz).

#### 2.3.7 Identification of outlier cases

All measurements where the Standard Deviation of NN/(RR) intervals ≥ 150 were visually inspected (n = 107). 13 of the identified measurements were suspected of measurement error (e.g. due to displaying doubling and then halving of IBI length). These were noted so that they could be excluded from the analysis.

#### 2.3.8 vmHRV indices of interest

Two commonly used indicators of vagal tone are the time domain measure RMSSD and the high-frequency component measure HFHRV (Task Force, 1996; Thayer et al., 2012). We were mainly interested in HFHRV due to RMSSD possibly being influenced by sympathetic activation (Berntson et al., 2005) and vagal (Huston et al., 2006; Wang H et al., 2003) and sympathetic (Madden et al., 1995; Vizi et al., 1998) inputs to the spleen having opposing effects on inflammatory marker expression. RMSSD and HFHRV are usually highly correlated (Goedhart et al., 2007). Due to the short data length of our recordings, RMSSD was included in the analysis to check for this correlation to assure that vmHRV indices were of good quality.

### 2.4 C Reactive Protein (CRP), Leukocyte (WBC) count, and Leukocyte telomere length (LTL) quantification

Blood sampling is described in Eggen et al. (2013). The participants were not required to fast, but they were only allowed to drink water and black coffee during their visits. 50 ml blood samples were collected (n = 12,882). Time since last meal (hours) was recorded. Venipuncture was performed with subjects in a sitting position. A light tourniquet was used and released before sampling. After 30 minutes at room temperature, the coagulated samples were centrifuged at 2000 g for 10 minutes, and the sera were transferred within 1 hour to plastic tubes, and kept between 1°C and 10°C. The blood samples were sent twice daily to the Department of Laboratory Medicine, University Hospital North Norway, Tromsø, which is an accredited laboratory (ISOstandard 17025).

CRP was analyzed by a highly sensitive CRP method (particle-enhanced immunoturbidimetric assay). The analyses were performed on a Modal PPE auto-analyzer with reagents from Roche Diagnostics Norway AS.

For WBC, 5 ml of blood was collected into Vacutainer tubes containing EDTA as an anticoagulant (K3-EDTA 40 ll, 0.37 mol/l per tube), and analyzed within 12 hr by an automated blood cell counter (Coulter CounterÒ, Coulter Electronics, Luton, UK and Coulter LH750, Nerliens Meszansky). Leukocyte telomere length (LTL) was analyzed by established qPCR methods (O’Callaghan & Fenech, 2011) at McGill University in 2018 and the mean telomere length was quantified as arbitrary units.

### 2.5 Vagal immunomodulation index

To calculate the neuroimmunomodulation (NIM) index, the high frequency component of HRV (HFHRV) was divided by participants’ CRP to obtain the HFHRV/CRP ratio in accordance with Gidron et al. (2018).

### 2.6 Confounding variables

One major issue facing the literature on associations between vmHRV and inflammatory markers is the lack of controlling for confounding variables (Haensel et al., 2008; Papaioannou et al., 2013). Timing from last meal to blood samples being drawn (Marinac et al., 2015), disturbed sleep (Irwin et al., 2018), depression (Lee et al., 2019), and alcohol and tobacco use (e.g. Tibuakuu et al., 2017; van de Loo et al., 2020) are all associated with inflammatory marker expression as well as PFC activity and vmHRV. Thus, measures assessing these variables were included as control variables in the analysis.

WBC has previously been found to be positively associated with LTL (Neuner et al., 2015) and telomere length varies between different subtypes of leukocytes (Rufer et al., 1998; Wong et al., 2011). Considering the positive associations between WBC and CRP (Sproston & Ashworth, 2018), and negative associations between CRP and LTL (Rode et al., 2014; Wong et al., 2014), WBC could serve as a positive confounder in the relationship between CRP and LTL. It could also serve as a confounder in the relationship between the vagal tone and LTL, as vagal and sympathetic activity differentially regulate leukocyte activity with vmHRV being negatively associated with WBC (Williams et al., 2019). WBC was therefore included as a control variable in the analysis.

### 2.7 Statistical analysis

First, visual inspection of the variables revealed that LTL, WBC, CRP, NIM index, and HRV were not normally distributed. Log10 transformation was performed on these variables prior to analysis. After log10 transformation, visual inspection confirmed normal distribution. All subsequent analyses were performed on the log10 transformed version of these variables.

Pearson correlations were conducted for HFHRV, RMSSD, LTL, the NIM index, CRP, and WBC to assess the relationship between the variables.

All regression analyses were checked for violation of assumptions regarding normality, linearity, and homoscedasticity by plotting standardized predicted values against standardized residuals, and multicollinearity was assessed by checking that variance inflation factor values ≈ 1. No assumptions were violated at any time. Mediation analyses were performed using a series of hierarchical multiple linear regression (two models per analysis). Due to a positive correlation between WBC and LTL, and WBC and CRP in our sample, WBC was used as a control variable in all regression analyses. In the first model of the first analysis, LTL was entered as the dependent variable and compared against the control variable, WBC, which was entered as an independent variable. Then, in the second model of the first analysis, HFHRV was entered as the independent variable and compared against the dependent variable to examine main effects of the HFHRV on the LTL (path C). In the first model of the second analysis, CRP was entered as the dependent variable with WBC as independent variable. In the second model of the second analysis, HFHRV was entered as the independent variable and compared against the dependent variable to examine main effects of the HFHRV on the CRP (path A). In the first model of the third analysis LTL was entered as the dependent variable and compared against the control variable, WBC, which was entered as an independent variable. Finally, in the second model of the third analysis, HFHRV and CRP were both entered as independent variables and compared against the dependent variable (LTL) to examine main effects of CRP on LTL (Path B) and HFHRV on the LTL (path C’) simultaneously. To calculate the total effect of path A and B on LTL, path A was multiplied with path B (A*B).

Multiple linear regression analysis was performed on the NIM index and LTL (NIM-LTL model). WBC was entered as the independent (control) variable and LTL as the dependent variable in model 1, and the NIM index and WBC as the independent variables and LTL as the dependent variable in model 2. When controlling for confounders, hierarchical multiple linear regression was used following the same steps as in the initial regression analysis. All control variables were individually entered as predictor variables in model 1 along with WBC, and with LTL as outcome variable. In the next step (model 2), HFHRV and CRP were entered simultaneously as predictor variables. This was repeated for all control variables. Regression analyses were also performed without WBC as a control variable to assess how its inclusion affected results. Control variables were either continuous or numerical.

Additional regression analyses were performed with a reduced dataset (n=169) excluding all participants that reported sleeping problems once per week or more during the past 12 months, symptoms of depression (feeling depressed, feeling worthless or useless, feeling hopelessness for the future), current or previous history of smoking or smokeless tobacco use, drinking more than 2-4 times per month, usually consuming more than four units of alcohol when drinking. This was done for both the mediation analysis and NIM index analysis while controlling for WBC. Descriptive statistics were generated for the reduced dataset. After exclusion of 13 outlier cases suspected of measurement error, regression analyses were repeated on the original (n = 1359) and reduced (n = 165) dataset.

Data was analyzed using SPSS version 25 (IBM Corp, 2017).

## 2 Results

### 3.1 Participant characteristics and descriptive statistics of vmHRV, NIM index, and biomarkers

The demographic characteristics of the studied population (N = 1372) are shown in Table 1. With respect to confounders of vmHRV, PFC, and inflammatory marker expression, 226 participants reported feeling depressed, 183 reported feeling useless or worthless, and 197 reported feeling hopelessness during the past week. 183 participants reported having sleeping problems more than once per week during the last 12 months. 280 participants reported alcohol use more than two days per week, 313 participants reported drinking more than 2 units of alcohol when drinking, while 26 participants reported drinking six or more units of alcohol once or more per week. 218 participants were daily smokers while 701 participants used to be daily smokers. 41 participants reported current use of snus/snuff/chewing tobacco. Table 2 provides descriptive statistics for extracted vmHRV and NIM indices, LTL, CRP, and WBC before and after log10 transformation, and time since last meal (hours).

**Table 2.** Descriptive statistics for vmHRV and NIM indices, Telomere length, CRP, Leukocyte count, and Time since last meal

| Variable | Mean | Std. | Minimum | Maximum | N |
| --- | --- | --- | --- | --- | --- |
| RMSSD | 32.98 | 40.38 | 4 | 563 | 1372 |
| RMSSD_log10 | 1.36 | .32 | .56 | 2.75 | 1372 |
| HFHRV | 300.37 | 1579.79 | 1 | 48970 | 1372 |
| HFHRV_log10 | 1.78 | .65 | -.11 | 4.69 | 1372 |
| NIM | 213.52 | 815.56 | .04 | 19203.73 | 1371 |
| NIM_log10 | 1.56 | .80 | -1.41 | 4.28 | 1371 |
| LTL | 1.31 | .57 | 0 | 5 | 1372 |
| LTL_log10 | .08 | .18 | -.62 | .69 | 1372 |
| CRP | 2.99 | 5.57 | 0 | 81 | 1371 |
| CRP_log10 | .22 | .42 | -.72 | 1.91 | 1371 |
| WBC | 6.3 | 2.07 | 1 | 39 | 1369 |
| WBC_log10 | .78 | .11 | .04 | 1.59 | 1369 |
| Time since last meal | 3.36 | 1.77 | 0 | 9 | 1364 |

*Notes.* RMSSD = Root mean square of successive differences. \_log10 = Log10 transformed variable. HFHRV = High frequency component heart rate variability. NIM = Neuroimmunomodulation index. LTL = Leukocyte telomere length arbitrary units. CRP = C-Reactive Protein. WBC = Leukocyte count.

### 3.2 Relationship between vmHRV, CRP, and LTL

RMSSD and HFHRV were highly correlated (r = .904, p < .001). Table 3 provides correlation between vmHRV indices, LTL, CRP, NIM index, and WBC. HFHRV was significantly associated with LTL (r = .068, p = .012) and CRP (r = -.068, p = .012). HFHRV was not significantly associated with WBC. The NIM index was significantly associated with LTL (r = .067, p = .013), CRP (r = -.579, p < .001), and WBC (r = -.113, p < .001). CRP was not significantly associated with LTL. CRP was significantly associated with WBC (r = .286, p < .001). WBC was significantly associated with LTL (r = .164, p < .001). Time since last meal was positively associated with LTL (r = .067, p = .014).

**Table 3.** Pearson correlations (r) between vmHRV, LTL, CRP, NIM, and WBC (N = 1372)

| Variables | 1 | 2 | 3 | 4 | 5 | 6 | 7 |
| --- | --- | --- | --- | --- | --- | --- | --- |
| 1 HFHRV | - |  |  |  |  |  |  |
| 2 RMSSD | .904*** | - |  |  |  |  |  |
| 3 NIM | .853*** | .760*** | - |  |  |  |  |
| 4 LTL | .068* | .047 | .067* | - |  |  |  |
| 5 CRP | -.068* | -.040 | -.579*** | -.018 | - |  |  |
| 6 WBC | .035 | .017 | -.113*** | .164*** | .286*** | - |  |
| 7 TSLM | .025 | .054* | .014 | .067* | .014 | .039 | - |

*Notes.* 2-tailed. All variables are log10 transformed. HFHRV = High frequency heart rate variability. RMSSD = Root means squared of successive differences. NIM = Neuroimmunomodulation index. LTL = Leukocyte telomere length arbitrary units. CRP = C reactive protein. WBC = Leukocyte count. TSLM = Time since last meal.
\* $p < .05$ . \*\* $p < .01$ . \*\*\* $p < .001$ .

Table 4 provides the results for the hierarchical regression model with HFHRV and CRP as predictor variables and LTL as outcome variable. In the first regression analysis, HFHRV was a significant positive predictor of LTL (β = .062, p = .020, *R^2^_adj_* = .026, *F* = 19.33) when controlling for WBC. In the second regression analysis, HFHRV was a significant negative predictor of CRP (β = -.077, p = .003, *R^2^_adj_* = .079, *F* = 59.42) when controlling for WBC. In the third regression analysis, HFHRV was a significant positive predictor of LTL, and CRP was a significant negative predictor of LTL (HFHRV: β = .057, p = .033; CRP: β = -.065, p = .020; *R^2^_adj_* = .029; *F* = 14.74) when controlling for WBC. The model with and without mediator is illustrated in Figure 3. CRP stopped being a significant predictor of LTL when not controlling for WBC.

**Table 4.** Multiple linear regression analysis for HFHRV, CRP, and LTL (N = 1372)

| LTL |  |  |  |
| --- | --- | --- | --- |
| Predictors (model 1) | $\beta$ | $R^2_{adj}$ | Sig. |
| WBC | .154 | .023 | .000 |
| Predictors (model 2) |  |  |  |
| WBC | .152 | .026 | .000 |
| HFHRV | .062 | .026 | .020 |
| CRP |  |  |  |
| Predictors (model 1) | $\beta$ | $R^2_{adj}$ | Sig. |
| WBC | .272 | .074 | .000 |
| Predictors (model 2) |  |  |  |
| WBC | .275 | .079 | .000 |
| HFHRV | -.077 | .079 | .003 |
| LTL |  |  |  |
| Predictors (model 1) | $\beta$ | $R^2_{adj}$ | Sig. |
| WBC | .154 | .023 | .000 |
| Predictors (model 2) |  |  |  |
| WBC | .170 | .029 | .000 |
| HFHRV | .057 | .029 | .033 |
| CRP | -.065 | .029 | .020 |

*Notes.* All variables are log10 transformed. All analyses controlled for leukocyte count. WBC = Leukocyte count. LTL = Leukocyte telomere length arbitrary units. HFHRV = High frequency heart rate variability. CRP = C reactive protein.

**Figure 3.**
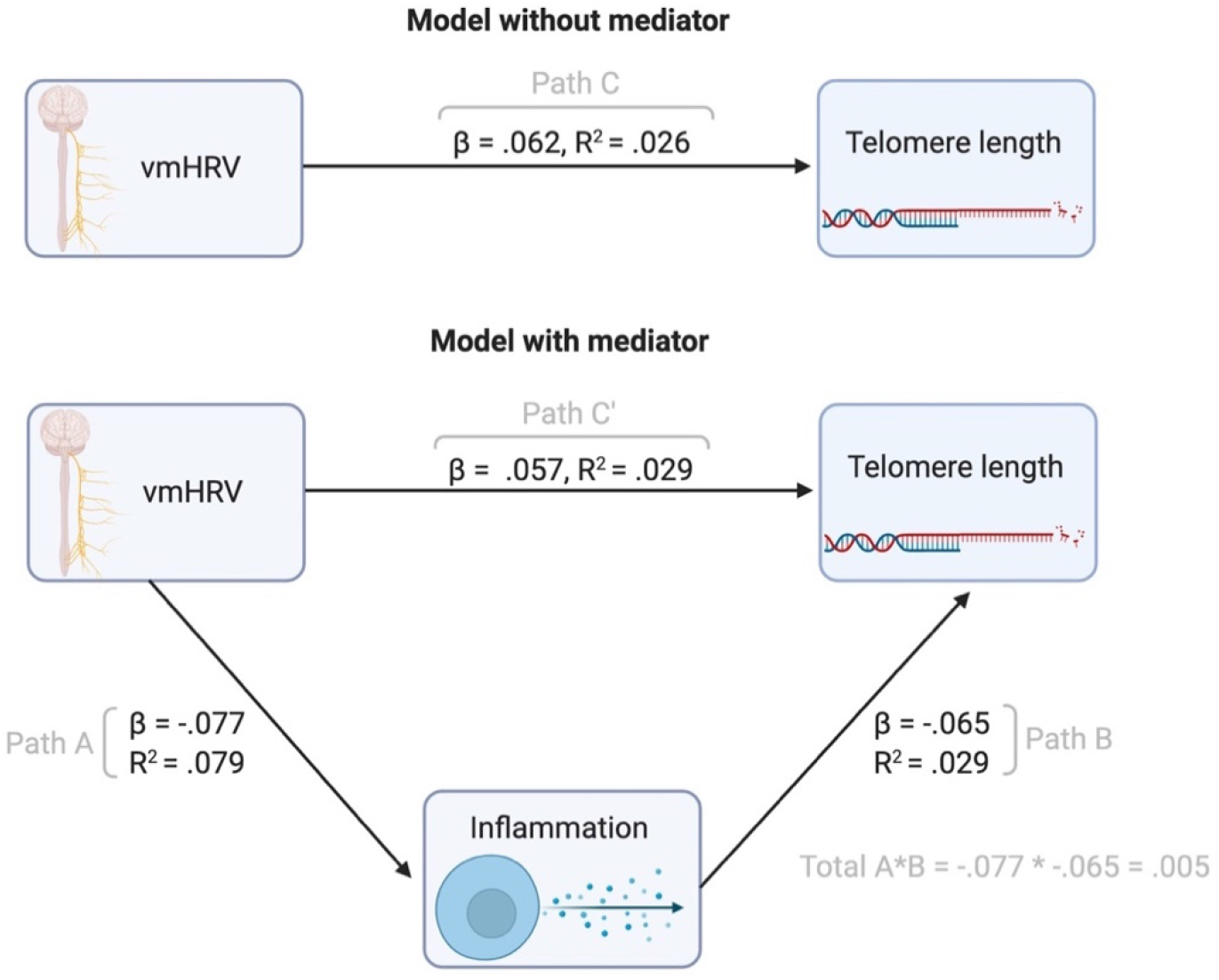
Regression model with and without mediator. All relationships controlling for leukocyte count. Figure created with BioRender.com.

**Figure 4.**
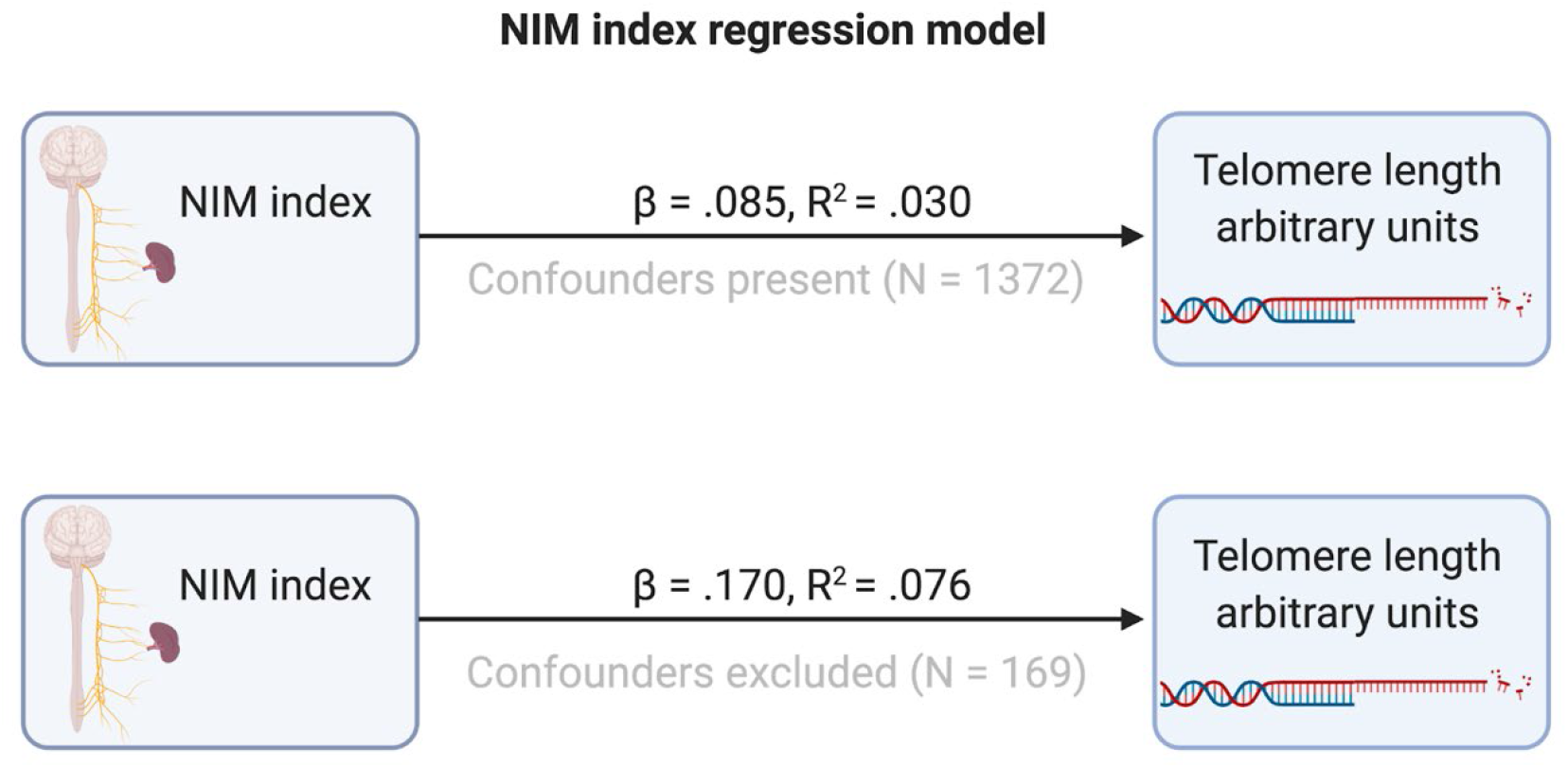
Regression model for the relationship between NIM index and telomere length. All relationships controlling for leukocyte count. Figure created with BioRender.com.

In the mediation model, all relationships remained significant when individually controlling for: time since last meal (HFHRV: β = .056, p = .035; CRP: β = -.066, p = .017; *R^2^_adj_* = .032, *F* = 12.21), sleeplessness during the last 12 months (HFHRV: β = .054, p = .048; CRP: β = -.067, p = .019; *R^2^_adj_* = .030, *F* = 11.13), frequency of alcohol consumption (HFHRV: β = .054, p = .045; CRP: β = -.067, p =.018; *R^2^_adj_* = .028, *F* = 10.78), and how often participants drink 6 units of alcohol or more on one occasion (HFHRV: β = .059, p = .038; CRP: β = -.057, p = .050; *R^2^_adj_* = .031, *F* = 10.78).

HFHRV did not remain a significant predictor of LTL when individually controlling for: sleeping problems during last week, feeling depressed during last week, feeling worthless or useless during last week, feeling hopelessness during last week, units of alcohol consumed when drinking, previous or current daily smoking, and age when smokers started smoking daily. CRP did not remain a significant predictor of LTL when independently controlled for: occasional smoking, years since smoking cessation, snuff/chewing tobacco use. Neither HFHRV nor CRP remained significant predictors of LTL when independently controlling for: number of cigarettes smoked per day, years of smoking.

After exclusion of 13 outlier cases (n = 1359) suspected of IBI measurement error (as described in section 2.3.7) when simultaneously assessing the effect of HFHRV and CRP on LTL, CRP but not HFHRV (CRP: β = -.066, p = .018; HFHRV: β = .052, p = .053; *R^2^_adj_* = .030, *F* = 15.04) remained a significant predictor of LTL when controlling for WBC.

### 3.3 Relationship between the NIM index and LTL

Table 5 provides the results for the hierarchical regression model with the NIM index as predictor variable and LTL as outcome variable. The NIM index was a significant positive predictor of LTL (β = .085, p = .001, *R^2^_adj_* = .030, *F* = 21.77) when controlling for WBC.

**Table 5.** Regression analysis for the NIM index and LTL (N = 1372)

| Predictors (model 1) | LTL |  |  |
| --- | --- | --- | --- |
| | $\beta$ | $R^2_{adj}$ | Sig. |
| WBC | .154 | .023 | .000 |
| **Predictors (model 2)** |  |  |  |
| WBC | .164 | .030 | .000 |
| NIM | .085 | .030 | .001 |

*Notes.* All variables are log10 transformed. All analyses controlled for leukocyte count. NIM = Neuroimmunomodulation index. WBC = Leukocyte count. LTL = Leukocyte telomere length, arbitrary units.

In the NIM-LTL regression model, the NIM index remained a significant predictor of LTL when individually controlling for: time since last meal (β = .086, p = .001, *R^2^_adj_* = .032, *F* = 16.02), sleeplessness during the last 12 months (β = .084, p = .002, *R^2^_adj_* = .030, *F* = 14.58), sleeping problems during last week (β = .082, p = .003, *R^2^_adj_* = .031, *F* = 14.81), feeling depressed during last week (β = .087, p = .002, *R^2^_adj_* = .033, *F* = 15.40), feeling worthless or useless during last week (β = .086, p = .002, *R^2^_adj_* = .031, *F* = 14.34), feeling hopelessness during last week (β = .087, p = .002, *R^2^_adj_* =.032, *F* = 15.07), frequency of alcohol consumption (β = .084, p = .002, *R^2^_adj_* = .028, *F* = 14.09), units of alcohol consumed when drinking (β = .078, p = .009, *R^2^_adj_* =.028, *F* = 11.73), how often participants drink 6 units of alcohol or more on one occasion (β = .083, p = .003, *R^2^_adj_* = .032, *F* = 14.36), occasional smoking (β = .084, p=.003, *R^2^_adj_* =.037, *F* = 16.32), previous or current daily smoking (β = .079, p = .003, *R^2^_adj_* = .036, *F* = 17.87), years since smoking cessation (β = .118, p = .002, *R^2^_adj_* =.028, *F* = 7.78), cigarettes per day (β = .081, p = .025, *R^2^_adj_* = .022, *F* = 6.65), age when smokers started smoking daily (β = .088, p = .009, *R^2^_adj_* = .023, *F* = 8.09), use of snuff or chewing tobacco (β = .076, p = .006, *R^2^_adj_* =.028, *F* = 14.44).

The NIM index did not remain a significant predictor of LTL when individually controlling for years of daily smoking. The NIM index remained a significant predictor of LTL (β = .067, p = .013, *R^2^_adj_* = .004, *F* = 6.20) when not controlling for WBC.

After exclusion of 13 outlier cases (N = 1359) suspected of IBI measurement error (as described in section 2.3.7), the NIM index was still a significant predictor of LTL (β = .083, p = .002, *R^2^_adj_* = .030, *F* = 22.30) when controlling for WBC.

### 3.4 Relationship between the vmHRV, CRP and LTL, and between NIM index and LTL in reduced dataset

To account for directionality and confounder effects on results, analyses where repeated where all participants where confounders were present were excluded (n = 169). Descriptive statistics for the reduced dataset can be found in Table 6.

**Table 6.** Descriptive statistics for vmHRV, NIM index, LTL, CRP, Leukocyte count, and Time since last meal in reduced dataset

| Variable | Mean | Std. | Minimum | Maximum | n |
| --- | --- | --- | --- | --- | --- |
| RMSSD | 37.81 | 58.88 | 4 | 563 | 169 |
| RMSSD_log10 | 1.38 | .35 | .60 | 2.75 | 169 |
| HFHRV | 654.51 | 3938.39 | 1 | 48970 | 169 |
| HFHRV_log10 | 1.83 | .73 | .17 | 4.69 | 169 |
| NIM | 354.88 | 1611.61 | .45 | 19203.73 | 169 |
| NIM_log10 | 1.63 | .82 | -.34 | 4.28 | 169 |
| LTL | 1.32 | .58 | 0 | 4 | 169 |
| LTL_log10 | .08 | .18 | -.45 | .65 | 169 |
| CRP | 2.62 | 4.36 | 0 | 38 | 169 |
| CRP_log10 | .20 | .38 | -.57 | 1.58 | 169 |
| WBC | 5.88 | 1.43 | 2 | 11 | 168 |
| WBC_log10 | .75 | .10 | .30 | 1.06 | 168 |
| Time since last meal | 3.27 | 1.62 | 0 | 9 | 169 |

*Notes.* RMSSD = Root mean square of successive differences. \_log10 = Log10 transformed variable. HFHRV = High frequency component heart rate variability. NIM = Neuroimmunomodulation index. LTL = Leukocyte telomere length arbitrary units. CRP = C-Reactive Protein. WBC = Leukocyte count.

When excluding all participants that reported sleeping problems, symptoms of depression, smoking or smokeless tobacco use, drinking more than 2-3 days per week, consuming more than four units of alcohol when drinking, and consuming six or more units of alcohol once or more per week, the NIM index (β = .170, p = .025, *R^2^_adj_* = .076, *F* = 7.84) remained a significant predictor of LTL, but not HFHRV nor CRP, when controlling for WBC. The NIM index remained a significant predictor of LTL when simultaneously controlling for time since last meal and WBC (β = .170, p = .026, *R^2^_adj_* =.070, *F* = 5.19).

After exclusion of 4 outlier cases (N = 165) suspected of IBI measurement error (as described in section 2.3.7), the NIM index was still a significant predictor of LTL (β = .162, p =.033, *R^2^_adj_* =.081, *F* = 8.17) when controlling for WBC. The NIM index remained a significant predictor of LTL when simultaneously controlling for time since last meal and WBC (β = .162, p = .035, *R^2^_adj_* =.075, *F* = 5.42).

## 4 Discussion

This is the first study to assess inflammation as a mediator in the relationship between vmHRV and telomere length, and to assess the relationship between the NIM index, a recently proposed index for vagal immunomodulation (Gidron et al., 2018) and possible indicator of PAS axis activity, and telomere length. It is also the first paper to conceptualize the neural substrate of these relationships anatomically as the PAS axis.

In the initial correlation analysis (Table 3), the effects of the positive association between vmHRV and telomere length were equal to the effect of the negative association between vmHRV and CRP. In our mediation analysis (Table 4), when controlling for WBC, we found that vmHRV had a direct positive effect on telomere length and a negative effect on CRP levels, while CRP had a negative effect on telomere length. These findings are in line with the hypothesis that inflammation is a negative mediator in the relationship between telomere length and vagal tone (Ask et al., 2018) and supports previous findings suggesting relationship between vmHRV and inflammation (Williams et al., 2019), vmHRV and telomere length (Perseguini et al., 2015; Streltsova et al., 2017; Woody et al., 2017; Zalli et al., 2013), and CRP and telomere length (Rode et al., 2014; Shin et al., 2019; Wong et al., 2014). The combined positive effects of vmHRV on CRP, and negative effects of CRP on telomere length via the indirect pathway almost cancel each other out, further suggesting that the balance between PAS axis activity and inflammatory output helps to preserve telomere integrity. Taken together, these mediation analyses provide partial statistical support for the hypothesis that inflammation mediates the relationship between vagal tone and telomere length as proposed by the NISIM (Ask et al., 2018).

Traditional mediation analyses have been critiqued for being too dependent on subjective interpretation (e.g. Agler & De Boeck, 2017). This could be a challenge in the present study, as vagal tone by itself is not a measure of inflammation, nor is CRP a measure of vagal tone. Evidence indicates that CRP is an important regulator of inflammation (Sproston & Ashworth, 2018), suggesting confounding effects that could partially de-couple its relationship with telomere length from PAS axis activity in the present study. The role of CRP in inflammation is not straightforward with sometimes opposing effects being ascribed to pentameric native CRP (nCRP) versus monomeric CRP (mCRP), the former isoform having primarily anti-inflammatory effects while mCRP having mainly pro-inflammatory effects (Sproston & Ashwort, 2018). In our study, CRP isoform was not specified, thus, the mediation analysis might therefore not provide sufficient sensitivity to properly examine the relationships of interest. One thing to keep in mind when using HRV indicators of vagal tone is that, while the sympathetic nervous system acts as a unit, the parasympathetic nervous system does not. Thus, there is arguably going to be some level of dissociation between vagal input to the heart and vagal input to the spleen. The recently proposed NIM index (Gidron et al., 2018) is based on the ratio of vmHRV to CRP (vmHRV/CRP) and could serve as an indicator of prefrontally modulated vagal immunomodulation thus arguably be a better proxy for the PAS axis. In our regression analysis, we found that the NIM index was a stronger positive predictor of telomere length than vmHRV.

A huge problem with studies on the relationship between inflammation and vagal tone is that they do not control for time since the last meal, sleeping habits and problems, depression, and alcohol and tobacco use (Haensel et al., 2008; Papaioannou et al., 2013). One strength of this study was that we controlled for these confounders. In our mediation analysis, all relationships remained significant for 4 of 18 confounding variables; vmHRV remained a significant positive predictor of telomere length for 7 out of 17 confounding variables, while CRP remained a significant negative predictor of telomere length for 11 out of 17 confounding variables. The NIM index remained a significant positive predictor of telomere length when controlling for 16 of 17 confounding variables. These results can be interpreted in several ways.

Depression is associated with lower PFC activity, and reduced PFC function is a suggested risk factor for development of depression (Hare & Duman, 2020; Kaya & McCabe, 2019). Tobacco use is associated with altered nicotinic acetylcholine receptor availability in the PFC (Brody et al., 2016; Feduccia et al., 2012) which are integral to its function, especially in the dorsolateral PFC (DLPFC; Croxson et al., 2011; Yang et al., 2013). This area of the PFC is one of the PFC components with the clearest relationships with vmHRV (Brunoni et al., 2013; Nikolin et al., 2017). Both chronic and acute alcohol use is associated with reduced PFC functioning (Abernathy et al., 2010; Olsen et al., 2015). The development and maintenance of addictive behaviors including smoking is associated with reduced PFC functioning (Olsen et al., 2015). Sleeping problems are also associated with reduced PFC functioning (e.g Vartanian et al., 2014). One possible interpretation could thus be that the confounders represent individuals with reduced PFC integrity that by consequence are more prone to the behaviors and conditions indicated by control variables, and that these are indeed the individuals for which the effects would be the most pronounced. Going by this interpretation, the effect of the confounders may not have implications on the significance of the results. An alternative interpretation could be owed to the inflammatory responses that these conditions and behaviors would produce that are not a direct consequence of (reduced) PFC input to the vagus nerve. For instance, oral nicotine administration decreases vmHRV in healthy non-smokers (Sjoberg & Saint, 2011), which, if independent of interactions with nicotinic receptors in the PFC could still have downstream effects on inflammatory marker expression. Inflammatory markers and ROS may have depressive effects on vmHRV, confounding the directionality of relationships (Lee MS et al., 2014). To account for these various interpretations, we did an additional regression analysis with a reduced dataset (from 1372 initial participants to 169) that excluded all participants where any of the confounders were present. In this dataset, only the NIM index remained a positive predictor of telomere length, and this relationship was significant after controlling for time since last meal. The explained variance and effect sizes in this reduced dataset were double of that which was observed in the initial dataset. The NIM index may therefore be a more sensitive proxy for the PAS axis and might mitigate some of the effects owed to lack of CRP isoform specification. After exclusion of some outlier cases suspected of measurement error, the NIM index remained a positive predictor of LTL in both the full and reduced dataset, indicating that this index is a robust indicator of PAS axis activity.

Due to the preliminary nature of the current findings and the indirect nature of variables that are used in the present study (and the possible implications this may have had for the reported effect sizes), it is premature to make any definite suggestions with regards to the clinical implications that the results may have. However, observations from the current literature on neurodegenerative diseases indicate PAS axis-associated relationships that are worthy of further exploration. The NISIM argues that PAS axis dysregulation will accelerate aging (Ask et al., 2018). The role of amyloid plaque in Alzheimer’s disease is undecided with accumulating evidence indicating that it might not be the pathogenic trigger driving the neurodegenerative cascade (e.g. Sturchio et al., 2021). Alternative explanatory mechanisms that trigger and interact with known antecedents to neurodegenerative pathology are thus needed. Studies on senescent cells in the brain may offer plausible causative factors for neurodegeneration (Baker et al., 2011; Chinta et al., 2015). Recent studies indicated that increases in peripheral inflammation is a prodromal indicator of dementia, with greater cognitive impairment being associated with increased levels of IL-6 and TNF-α in Alzheimer’s and Parkinson (King et al., 2017; Wood, 2018; see Maletic & Raison (2014) for a review of how peripheral inflammatory signals can be transported into the brain in psychiatric disease). This may indicate that PAS axis-targeted interventions may protect against the development of age related disease in vulnerable populations.

While the effects proposed by the NISIM are small and accumulates over a lifetime in healthy populations (Ask et al., 2018), evidence from studies on cancer patients (Chiang et al., 2013; De Couck et al., 2013; Mouton et al., 2010, 2011, 2012), sepsis patients (de Castilho et al., 2018) and schizophrenic patients (Benjamin et al., 2020; North et al., 2021; Yu et al., 2008) indicate that the consequences of PAS axis dysfunction might be more severe in pathological conditions characterized by reduced prefrontally modulated vagal tone. If further validation of the NISIM is achieved, the findings in the above-mentioned clinical conditions may suggest avenues for neuroimmunomodulation treatments targeting the different levels of the PAS axis: (1) the level of the PFC, (2) the level of the ANS, (3) the level of the spleen and leukocytes, and at (4) the cognitive/behavioral level.

Future studies should determine the genes relevant for the development of the PFC’s ability to modulate vagal tone such that regenerative genetic interventions for increasing the PFC’s capacity for vagal modulation can be developed.

### 4.1 Limitations and future perspectives

The findings detailed in the present study have small effect sizes. Although the effects proposed by the NISIM *are small* and accumulate over the course of a lifetime (Ask et al., 2018), it is possible that some of the results can be attributed to the specific limitations of the current study. CRP may have opposing effects on inflammation depending on subtype (Sproston & Ashwort, 2018) and transcriptional induction of CRP is downstream of IL-6 (Boras et al., 2014). It is thus hard to say how the results would have been if CRP type had been specified or if IL-6 had been assessed instead. The present study should be repeated with CRP subtype specified as well as for other inflammatory markers, particularly IL-6 as this marker may provide a better proxy for the inflammatory effects of a dysregulated PAS axis (Williams et al., 2019). Another limitation is that the recordings used to quantify vmHRV were 30 seconds long which is short. The recording should last for at least 10 times the wavelength of the lower frequency bound of the investigated component, and 5 minute recordings is the preferred standard. It has, however, been shown that useful frequency-domain information can be derived from recordings as low as 40 s for HFHRV (Salahuddin et al., 2007). The RMSSD and HFHRV indices were highly correlated in our study, which indicate that the measures were of good quality. Future studies should use 5 minute recordings to quantify vmHRV to reduce effects owing to suboptimal IBI measurements. A third limitation owes to the correlational nature of the study as it does not provide direct evidence for the NISIM. Evidence suggests that telomere dysfunction and senescence can occur irrespective of telomere length (Victorelli & Passos et al., 2017). A recent study demonstrated that ROS-induced telomere dysfunction results in senescence by replication fork arrest (Coluzzi et al., 2019). Longitudinal studies are needed to assess the directional relationship between prefrontally modulated vagal tone, inflammatory markers, and telomere length, and should include markers of hydrogen peroxide induced telomere damage such as 8-oxoguanine, indicators of replication fork stall such as γH2AX with respect to 53BP1 at telomeres, and increases in the H3K9me3 histone mark (Coluzzi et al., 2019). Assessing the relationship between indicators of PFC functioning and markers of oxidative DNA damage is important to disentangle directionality problems such as telomere dysfunction preceding reduced prefrontal function or vice versa (e.g. Powell et al., 2018). A final limitation of the present study is that we were not interested in the effects of sex and did therefore not include this variable in the dataset and analysis. There have been reports of sex differences in vmHRV (e.g. Koenig & Thayer, 2016) and Alzheimer’s impact and progression (Laws et al., 2018), thus, future studies should examine sex differences in PAS axis dysregulation and its clinical relevance.

### 4.1 Conclusion

This study is an important first step towards understanding the anatomical-to-molecular mechanisms that relate individual differences in neurocognitive stress regulation capacity to PAS axis-related induction of cellular senescence and age-related pathology. Our results indicate that inflammation is downstream of PAS axis activity, suggesting that PAS axis dysregulation leads to increased levels of inflammation and subsequently ROS induced telomere damage. Based on the findings of our study, we argue that the NIM index is a sensitive proxy for PAS axis activity and could serve as a useful indicator when assessing PAS axis-related effects on antecedents to cellular senescence. Future studies should assess the relationship between indicators of prefrontally modulated vagal tone, IL-6, CRP subtype, and telomere length as well as more direct markers of oxidative telomere damage and telomere damage-induced cellular senescence.

## 5 Acknowledgements

We would like to thank our colleagues Leiv Arne Rosseland, Naomi Azulay, and Christian Tronstad for sharing their expertise and valuable comments that greatly improved the manuscript. We would particularly like to thank Leiv Arne Rosseland for help with organizing the study, Christian Tronstad for analysis of IBIs and quantification of HRV indices, and Naomi Azulay for help with detecting outliers in the HRV data.

## Notes

### Competing Interest Statement

The authors have declared no competing interest.

